# Genetic Substrates of Brain Vulnerability and Resilience in Aging APOE2 Mice

**DOI:** 10.1101/2022.12.12.520146

**Authors:** Ali Mahzarnia, Hae Sol Moon, Jacques Stout, Robert J Anderson, Madison Strain, Jessica T. Tremblay, Zay Yar Han, Andrei Niculescu, Anna MacFarlane, Jasmine King, Allison Ashley-Koch, Darin Clark, Michael W Lutz, Alexandra Badea

## Abstract

Understanding the interplay between genotype, age, and sex has potential to reveal factors that determine the switch between successful and pathological aging. APOE allelic variation modulate brain vulnerability and cognitive resilience during aging and Alzheimer disease (AD). The APOE4 allele confers the most risk and has been extensively studied with respect to the control APOE3 allele. The APOE2 allele has been less studied, and the mechanisms by which it confers cognitive resilience and neuroprotection remain largely unknown. Using mouse models with targeted replacement of the murine APOE gene with the human major APOE2 alleles we sought to identify changes during a critical period of middle to old age transition, in a mouse model of resilience to AD. Age but not female sex was important in modulating learning and memory estimates based on Morris water maze metrics. A small but significant 3% global brain atrophy due to aging was reflected by regional atrophy in the cingulate cortex 24, fornix and hippocampal commissure (>9%). Females had larger regional volumes relative to males for the bed nucleus of stria terminalis, subbrachial nucleus, postsubiculum (~10%), and claustrum (>5%), while males had larger volumes for the orbitofrontal cortex, frontal association cortex, and the longitudinal fasciculus of pons (>9%). Age promoted atrophy in both white (anterior commissure, corpus callosum, etc.), and gray matter, in particular the olfactory cortex, frontal association area 3, thalamus, hippocampus and cerebellum. A negative age by sex interaction was noted for the olfactory areas, piriform cortex, amygdala, ventral hippocampus, entorhinal cortex, and cerebellum, suggesting faster decline in females. Fractional anisotropy indicated an advantage for younger females for the cingulate cortex, insula, dorsal thalamus, ventral hippocampus, amygdala, visual and entorhinal cortex, and cerebellum, but there was faster decline with age. Interestingly white matter tracts were largely spared in females during aging. We used vertex screening to find associations between connectome and traits such as age and sex, and sparse multiple canonical correlation analysis to integrate our analyses over connectomes, traits, and RNA-seq. Brain subgraphs favored in males included the secondary motor cortex and superior cerebellar peduncle, while those for females included hippocampus and primary somatosensory cortex. Age related connectivity loss affected the hippocampus and primary somatosensory cortex. We validated these subgraphs using neural networks, showing increased accuracy for sex prediction from 81.9% when using the whole connectome as a predictor, to 94.28% when using the subgraphs estimated through vertex screening. Transcriptomic analyses revealed the largest fold change (FC) for age related genes was for Cpt1c (log2FC = 7.1), involved in transport of long-chain fatty acids into mitochondria and neuronal oxidative metabolism. Arg1, a critical regulator of innate and adaptive immune responses (log2FC = 4.9) also showed age specific changes. Amongst the sex related genes, the largest FC were observed for Maoa (log2FC = 4.9) involved in the degradation of the neurotransmitters serotonin, epinephrine, norepinephrine, and dopamine, and implicated in response to stress. Four genes were common for age and sex related vulnerability: Myo1e (log2FC = −1.5), Creld2 (log2FC = 1.4), Ptprt (log2FC = 2.9), and Pex1 (log2FC = 3.6). We tested whether blood gene expression help track phenotype changes with age and sex. Genes with the highest weight after connectome filtering included Ankzfp1 with a role in maintaining mitochondrial integrity under stress, as well as Pex1, Cep250, Nat14, Arg1, and Rangrf. Connectome filtered genes pointed to pathways relate to stress response, transport, and metabolic processes. Our modeling approaches using sparse canonical correlation analysis help relate quantitative traits to vulnerable brain networks, and blood markers for biological processes. Our study shows the APOE2 impact on neurocognition, brain networks, and biological pathways during a critical middle to old age transition in an animal model of resilience. Identifying changes in vulnerable brain and gene networks and markers of resilience may help reveal targets for therapies that support successful aging.

## Introduction

It is important to identify risk mitigating strategies to age associated decline, before irreversible brain damage caused by pathology and neurodegeneration. Prevalent amongst neurodegenerative diseases, Alzheimer’s disease (AD) affects ~10% of population aged 65 and the percentage of affected population increases with age. The major genetic risk for AD is related to the allelic variations of the APOE gene, a lipid transporter gene. The APOE4 allele confers the most risk and has been intensely studied with respect to the control APOE3 allele. The APOE2 variant promotes longevity and is neuroprotective, reducing risk against AD by nearly 50% (Conejero-Goldberg, Gomar et al. 2014), yet this allele has been less studied, and the mechanisms by which it confers cognitive resilience and neuroprotection remain largely unknown.

To understand the mechanisms through which APOE2 modulates biological pathways to protect cognition and confer brain resilience we need to develop methods that integrate over multivariate data sets. Relating cognition, imaging, and omics data has the potential to reveal image markers for resilience versus risk for cognitive decline and underlying biological pathways. GWAS studies (Harold, Abraham et al. 2009, Ballard, Gauthier et al. 2011, Lambert, Ibrahim-Verbaas et al. 2013) (Saykin, Shen et al. 2015) have revealed multiple genes that increase risk for pathological brain aging. To understand how aging influences brain network integrity, and reveal associated gene expression changes, researchers have traditionally relied on access to brain tissue. While brain based transcriptomic studies can provide powerful insight into gene expression changes and have sensitivity to location and cell types, such samples are rarely available from younger individuals. Brain samples for normal or cognitively impaired individuals undergoing life transitions are scarce, while MRI and peripheral markers, e.g., blood provide easily accessible data. More recently, blood-based RNA-seq data has been shown to relate to brain based gene interactions and processes (Panitch, Hu et al. 2022). Novel blood based RNA-seq analyses have also related gene expression levels to memory function (Niculescu, Le-Niculescu et al. 2020). It remains to be seen how these relationships map onto brain circuits, to help understand the role of APOE2 in conferring protection to cognitive decline during aging.

Mouse models replicating the human APOE alleles through targeted replacement of the murine APOE gene provide useful tools to address questions on the role of APOE genes in aging. We aimed to identify changes during middle to old age transition, and to examine the role of female sex during this time, in a model of successful aging and neuroprotection conferred by the APOE2 genotype (Conejero-Goldberg, Gomar et al. 2014). We focused on the transition between middle and old age, dichotomized into a young and old category, because of its significance with respect to the age of onset for neurodegenerative diseases, and the opportunities presented for implementing interventions if we can detect subtle biomarker changes at prodromal stages, before cognitive decline typical of AD.

To evaluate cognition during this critical transition period we have used the Morris water maze, to which we have recently introduced a novel metric (Badea, Li et al. 2022), the absolute winding number to complement learning and memory markers provided with an estimate of spatial navigation strategy. We used diffusion MRI to reveal atrophy and microstructural changes during this period and supplemented widely used measures of tissue integrity from fractional anisotropy with estimates of cellularity from MAP MRI. MAP MRI expands the diffusion MR signal in the local DTI reference frame using a complete set of orthogonal basis functions related to the eigenfunctions of the Fourier transform. Because of the dual nature of these basis functions, the measured signal in **q**-space and the propagator in **r**-space are represented with the same series coefficients, resulting in increased robustness to noise and signal confounds relative to traditional diffusion metrics (Avram, Sarlls et al. 2016). We estimated cellularity and restricted diffusion (Özarslan, Koay et al. 2013, Avram, Sarlls et al. 2016) in tissue based on the return-to-origin probability (RTOP). We hypothesized that such microstructural changes detected by MRI are accompanied by gene expression changes in the brain. But brain and even cerebrospinal fluid markers are difficult to obtain relative to blood biomarkers, who hold a higher potential for applicability to experimental studies of human aging process or neurodegenerative disease. Thus, there is interest in examining brain related changes from the perspective of blood biomarkers.

Recently the fusion of imaging and omics data has received increased attention. Here we propose modeling approaches that integrate through canonical correlation omics data from blood based transcriptomics, imaging data from brain MRI and derived connectomes describing networks, to memory related traits. These models seek to identify blood gene expression markers that help track brain and behavior phenotypes. If successful, such models will help understand the biological mechanisms that confer vulnerability or resilience during aging, and how genotype and aging interact to differentially affect male and female sex. Our study reveals how APOE2 impacts neurocognition, and modulates brain networks, and associated biological pathways changing during aging. More APOE2 studies can help inform the design of future strategies for enhancing the chances for successful aging and resilience to AD.

## Methods

### Animals

We used mice homozygous for the human APOE2 alleles (Sullivan, Mezdour et al. 1998) (targeted replacement), thought to be protective against Alzheimer’s disease. Our groups included 13 mice (8 males, 5 females) aged at the beginning of behavior experiments to ~12 months (12.64 ±0.70 months), and 28 mice aged to ~16 months (16.42±0.72) to examine age and sex specific differences during the transition from the human equivalent of middle age (10-14 months in a mouse, corresponding to 38-47 years on human), to old age (18-24 months in a mouse, corresponding to 56-69 years old in a human) (https://www.jax.org/news-and-insights/jax-blog/2017/november/when-are-mice-considered-old). At the end of behavior experiments we prepared brain specimens for imaging. Animals were anesthetized to a surgical plane using a mixture of ketamine/xylazine (100 mg/Kg ketamine, 5-10 mg/Kg xylazine), and perfused through the left cardiac ventricle, with outflow from the right atrium. Blood was flushed out with Saline (0.9%), at a rate of 8 ml/min, for ~5 min. For fixation we used a 10% solution of neutral buffered formalin phosphate (PBS) containing 10% (50 mM) Gadoteridol (ProHance, Bracco Diagnostics Inc., Monroe Township, NJ, United States), at a rate of 8 ml/min for ~5 min. Specimens were fixed in formalin for 12 hours then transferred in PBS plus 0.05% (2.5 mM) Gadoteridol until scanning in fomblin.

### Behavior

We tested spatial memory and learning through the Morris Water Maze test, as in (Badea, Wu et al. 2019) (Badea, Li et al. 2022). Mice were handled for 5 days prior to behavioral testing to habituate them to the researchers, and to water. Mice were then trained for 5 days in a 150 cm diameter circular swimming pool, where the water was rendered opaque using non-toxic white paint. Swim paths and times were tracked with a ceiling-mounted video camera, using ANY-maze (Stoelting, Wood Dale, IL, United States) software. Four trials were administered each day, in blocks of 2, separated by 30 min, and trials ended after 1 min maximum. Each trial consisted of placing the mouse into the water at one of four different starting positions, one in each quadrant. The quadrant order was varied each day. Mice could use visual cues to help find a platform submerged ~1.5 cm underneath the water. Mice were expected to learn that the platform is located in the same position using the visual cues, and locate it more quickly over time. We assessed learning by measuring the distance mice needed to swim to reach the platform, the percent swim distance in the target quadrant in which the platform was located, and the shape of the swim trajectory. If mice were unable to locate the platform within the allotted time of 1 min, they were guided to the platform and allowed to remain there for 10 s. Probe trials were conducted on days 5, 1 h after the last training trial, and then on day 8. During the probe trials the submerged platform was removed and mice were given 1 min to swim in the pool. Navigation strategies and efficiency were assessed using the total swim distance, the distance spent in the target quadrants, and the absolute winding number (Badea, Li et al. 2022). Animals were sacrificed ~ 2 months after the end of behavior testing.

### Imaging

Mouse brain specimens were imaged at 9.4T, as described in (Badea, Wu et al. 2019). We used a 3D spin echo diffusion weighted imaging (SE DWI) sequence with TR/TE: 100 ms/14.2 ms; matrix: 420 × 256 × 256; FOV: 18.9 mm × 11.5 mm × 11.5 mm, BW 62.5 kHz; reconstructed at 45 μm isotropic resolution. Diffusion weighting was applied along 46 directions, using 2 diffusion shells (23 at 2,000, and 23 at 4,000 s/mm^2^); and we acquired 5 non-diffusion weighted images (b0). The max diffusion pulse amplitude was 130.57 Gauss/cm; duration 4 ms; separation 6 ms. We used eight fold compressed-sensing acceleration (Uecker M; Ong F; Tamir JI; Bahri 2015) (Anderson, Wang et al. 2018) (Wang, Anderson et al. 2018). Diffusion and SHORE parameters were reconstructed using DIPY (Garyfallidis, Brett et al. 2014) with Q-ball Constant Solid Angle Reconstruction, producing ~2 million tracts. We have used pipelines implemented in a high-performance computing environment for voxel based analyses, and to calculate connectomes (Anderson, Cook et al. 2019), with reference to a symmetrized mouse brain atlas (Calabrese, Badea et al. 2015). Connectomes were normalized so that the diagonal was removed then the connectome and elements were divided by the sum of remaining entries.

### Transcriptomics

RNA-Seq experiments were carried out at the Duke Sequencing and Genomic Technologies Shared Resource Core, to identify differentially expressed genes with age, and sex. RNA was extracted from whole blood using QIAGEN kits, and we performed quality assurance using a NanoDrop 2000 spectrophotometer. The samples were processed using an Illumina NovaSeq600 S2 platform, 50 bp PE full flow cell, and NuGEN mRNA-Seq with Any Deplete Globin. Sequencing data were subjected to quality control using FastQC, using a cutoff threshold Phred > 20. There were no adapter sequences found flanking transcript reads, so data trimming was not performed and the raw data was used in the subsequent alignment and quantification steps. Raw RNAseq reads were aligned to a reference transcriptome (Mus_musculus.GRCm39.cdna.all.fa.gz) and the relative abundance of each transcript was quantified with the Salmon program (Patro, Duggal et al. 2017). A principal component analysis was done to identify confounding metadata factors that may influence the differential expression analysis (batch, qubit). The DESeq2 program (Love, Huber et al. 2014) was used for normalization and to assess differences in gene expression. Genes were annotated with their corresponding human gene name (ortholog for mouse) using BioMart (Durinck, Moreau et al. 2005). Gene set enrichment analysis (Subramanian, Tamayo et al. 2005) (Mootha, Lindgren et al. 2003) was performed to identify gene ontology terms and pathways associated with altered gene expression using https://tools.dice-database.org/GOnet/ (Pomaznoy, Ha et al. 2018).

### Statistical Analyses and Modeling

#### Behavior

We used R and the packages lme4, 1.1–27.1, and lmertest 3.1–3 to build mixed effect models for the learning trials with fixed effects for age, sex and time (Stage), and random effects for animal identity, e.g., Distance~Age*Sex*Stage+(1| AnimalID). We used linear models for the probes, e.g., Distance~Age*Sex. We used ANOVA to determine the effects of genotype and sex on behavioral markers including the total swim distance, normalized swim distance in the target quadrant, and the absolute winding number. ANOVA was followed by *post hoc* tests (using Sidak adjustments), and *p*< 0.05 was considered significant. Cohen’s F effect sizes were estimated using the package effsize 0.8.1.

#### Imaging

Regional and voxel wise analyses were conducted as in (Badea, Wu et al. 2019). The Statistical Parametric Mapping SPM toolbox (Friston, Worsley et al. 1994), version 12 was used with cluster false discovery rate correction.

#### Integrative Modeling

To achieve interpretable results from connectome data, we used two different methods that find the association between two types of data, e.g. connectome and traits (vertex screening), and between multiple data sets, e.g. connectomes, traits, and RNA sequencing data (sparse multiple canonical correlation analysis).

The <u>*vertex screening*</u> method identifies a signal subgraph by gradually removing nodes and computing the maximum distance correlation between the connectivity values of the remaining subgraph and the traits (Wang, Shen et al. 2018). We estimated the distance between graphs associated with different traits after iteratively removing the bottom 5% of nodes, iterating until the 5% of nodes were left. The final output is a set of vertices, and the maximum distance correlation between their connectivity and the traits. To validate this method, we constructed two Forward Neural Network (NN) models, one predicting the trait based on the extracted subgraph, and the other based on the full connectome matrices. We vectorized the full connectomes and filtered them using vertex screening, then estimated the average out of sample accuracy through k-fold cross-validation (k=5) using a multilayer perceptron NN and a single layer (logistic). We also implemented graph neural networks based on BrainGNN (Li, Zhou et al. 2021), and optimized the method for structural connectomes in our mouse models. The connectomes provided input to a graph convolutional network (GCN) layer followed by a topK pooling layer (Gao and Ji 2022), with another GCN layer and a topK pooling layer followed by two batch normalization and dropout layers and finally a softmax layer for classification. The GCN layers consist of ReLU activation function and the topK pooling layers consist of sigmoid activation function. We estimated prediction accuracy through a 5-fold strategy using an 80:20 data split.

The <u>*Sparse Multiple Canonical Correlation Analysis (SMCCA)*</u> method (Witten and Tibshirani 2009) seeks sparse canonical variables such that the linear combinations of three or more multivariate variables maximize the sum of their two-by-two correlations:

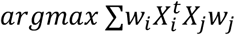

Subject to 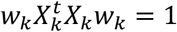 for all k.

This method results in a sparse and interpretable set of coefficients associated with connectomes. Feeding the model with RNA-seq data, traits, and winding numbers as the other random variables produces an interpretable set of panelized coefficients. To validate this model, we used 1000 bootstrap re-sampling with replacement to construct the 95^th^ percentile confidence interval for the sum of three correlations.

To understand the enriched processes for differentially expressed genes that were associated with changes in connectivity we used GOnet (Pomaznoy, Ha et al. 2018), available from https://tools.dice-database.org/GOnet/.

Our general approach on modeling age and sex associated vulnerability and resilience is described in **Figure 1**.

**Figure 1.**
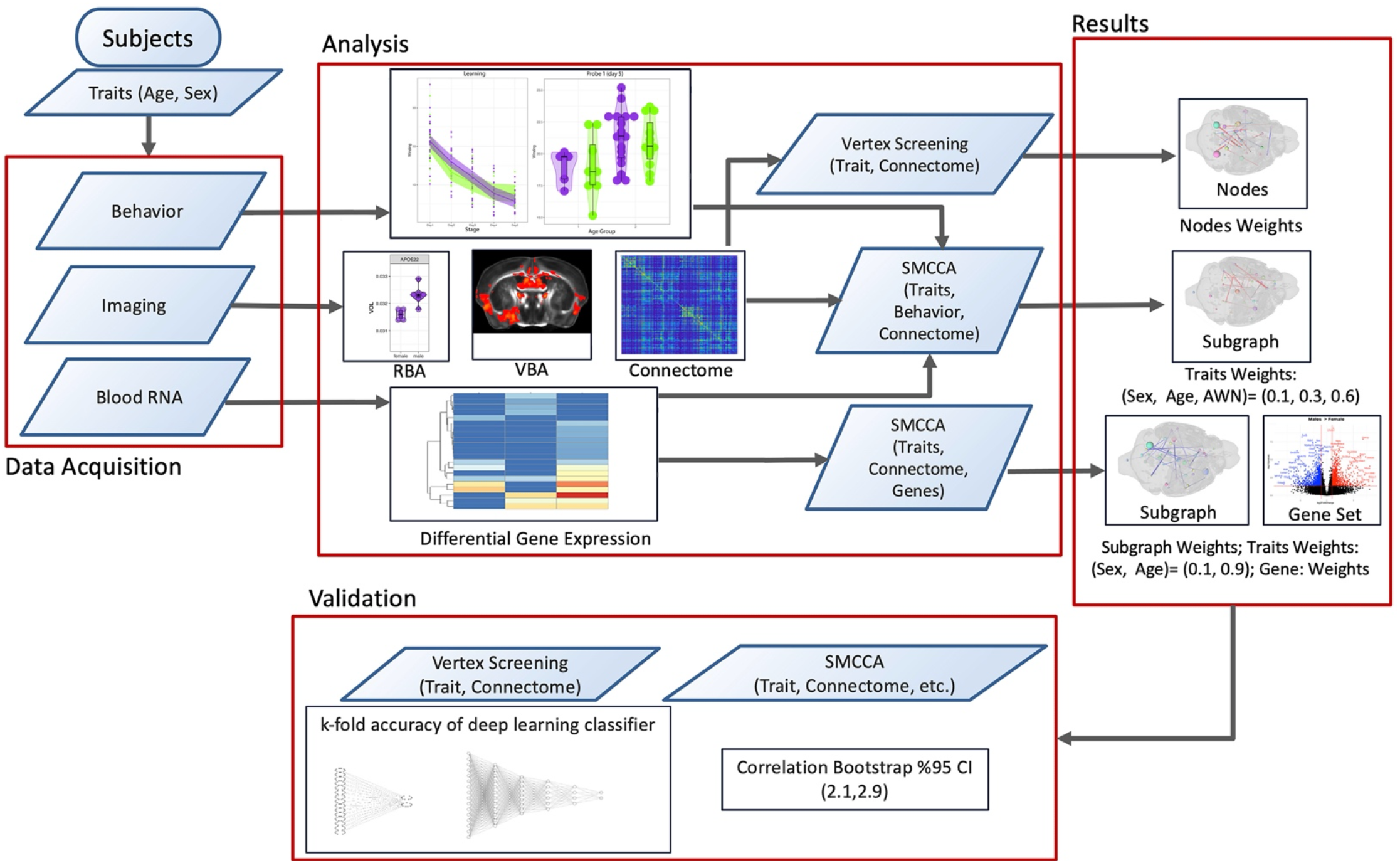
Schematic of the approach. To reveal brain and gene networks underlying differences with sex and aging we have acquired data on behavior, imaging, and blood RNA-seq which were analyzed individually, and using integrative modeling approaches such as vertex screening and sparse multi CCA (SMCCA). These were validated using bootstrap confidence intervals and k fold accuracy testing after a neural network based classifier.

#### Data and Code Availability Statement

Code and data necessary to reproduce the original analyses are available from https://github.com/AD-Decode/APOE2_Mouse.

#### Ethics Statement

All animal procedures have been approved by the Duke University IACUC committee.

## Results

To better understand the role of human APOE polymorphism, and in particular the mechanisms through which APOE2 confers resilience in aging and AD we have tested the dynamics of mouse brain networks in relation to cognitive and gene expression changes during the transition between middle and old age. Our results have revealed sex specific alterations in brain networks associated with memory function, and gene expression changes during aging.

### Memory changes in aging APOE2 mice

We have examined learning and memory using a Morris Water Maze test in mice aged to 12 and 16 months; (12.64±0.70 months; 8 males and 5 females), and (16.42±0.72 months; 11 males and 17 females) (**Figure 2**). Our estimates of learning and memory function are based on the distance mice swam in the pool until finding a submerged platform, the percent distance swam in the target quadrant containing the platform, and the absolute winding number characterizing the shape of the swim trajectories.

**Figure 2.**
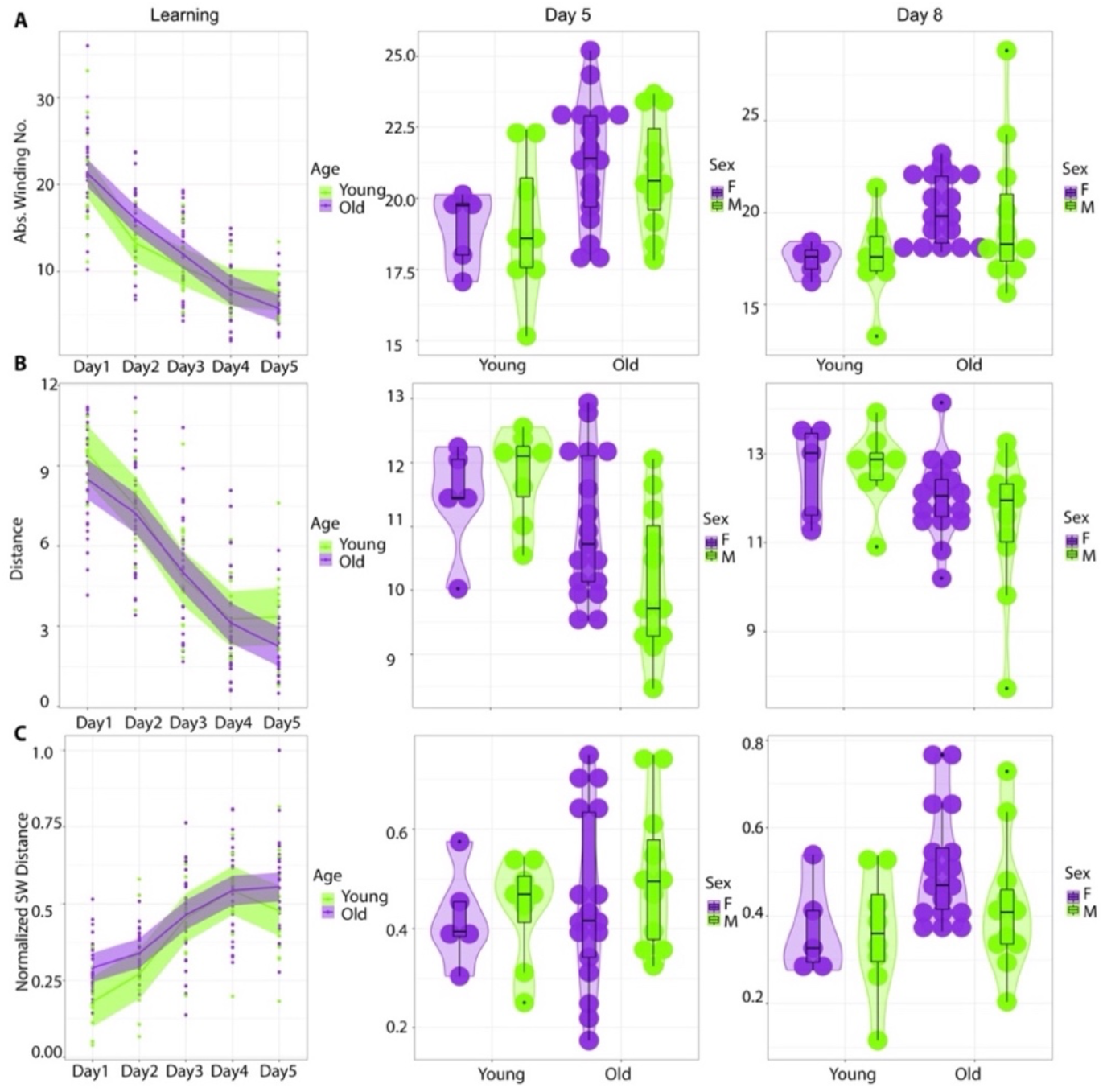
Learning and memory in APOE2 mice transitioning from middle to old age. We detected significant age but not sex effects for learning and memory based on the Morris water maze test. A. There was a significant interaction of age x stage for the absolute winding number during the learning trials. The winding number increased with age, indicating more complex trajectories in old relative to younger mice. B. The total distance did not change during the learning trials but decreased in day 5, and day 8 probes. C. The normalized distance in the target quadrant during learning trials and first probe was similar for the two groups, but increased in aged mice for the second probe.

For the learning trials sex and age differences were not significant, however we detected an effect of time in all three metrics (distance, normalized SW distance, winding number). For the total distance we obtained F(4,763) = 6.74; p = 2.49e-05; Cohen F = 0.19; CI = [0.10, 0.25]. For normalized distances we obtained F(4, 758.32) = 7.59, p = 5.32e-06; Cohen F = 0.20, CI = [0.12, 0.26]. For the absolute winding number we obtained F(4, 763) = 5.86, p = 0.0001, Cohen F = 0.18, CI = [0.09, 0.24]. There was a significant interaction of age x stage for the absolute winding number only: F(4, 763) = 2.85 p = 0.02, Cohen F = 0.12, CI = [0.01, 0.18], supporting increased sensitivity of the winding number to capture age differences.

For the first probe distance, age was significant (F1,37) = 8.36, p = 0.0064, Cohen F = 0.48, CI = [0.13, 0.81], and age x sex interaction was characterized by p = 0.1. There was no significant effect for the normalized target distance. Age was significant for the winding number F(1,37) = 8.93, p=0.005, Cohen F = 0.49, CI = [0.15, 0.83].

For the second probe there was a significant age effect for distance F(1,37) = 5.03, p = 0.03, Cohen F = 0.37, CI = [0.00, 0.70]; normalized distance in the target quadrant F = 5.95, p = 0.02, Cohen F = 0.40, CI = [0.05, 0.73], and the absolute winding number F(1,37) = 8.39, p = 0.006, Cohen F = 0.48, CI = [0.13, 0.81]. Our results support that age was an important factor modulating learning and memory, while female sex was not a detectable indicator for vulnerability.

### Image based phenotypes in aging APOE2 mice

#### Morphometry

The total brain volumes did not differ by sex, but declined from 527.91±9.86 mm^3^ to 512.35±17.41 mm^3^; this small 3% change was significant F(1,28) = 7.47, p = 0.01. A regional analysis revealed that females had larger volumes relative to males for the bed nucleus of stria terminalis, subbrachial nucleus, postsubiculum (~10%), and claustrum (>5%); while males had larger volumes for the brachium of the superior colliculus, orbitofrontal cortex, frontal association cortex, and the longitudinal fasciculus of pons >9%). The largest age associated atrophy was observed for the ventral hippocampal commissure (15%), fornix, and cingulate cortex areas 24 (9%). Opposite effects were noted for the parietal cortex, parietal association cortex, primary somatosensory cortex ~18% (**Supplementary Table 1**).

Voxel-based analysis (VBA), (**Figure 3**) revealed the spatial distribution of atrophy associated with sex and age. Compared to females, males had larger olfactory areas, dorsal caudate, piriform and cingulate cortices, preoptic areas, M1, amygdala, hippocampus, hypothalamus, and visual cortex, as well as a larger left cerebellum. In contrast females had larger clusters for the right caudate, hippocampus, entorhinal cortex, and right cerebellum.

**Figure 3.**
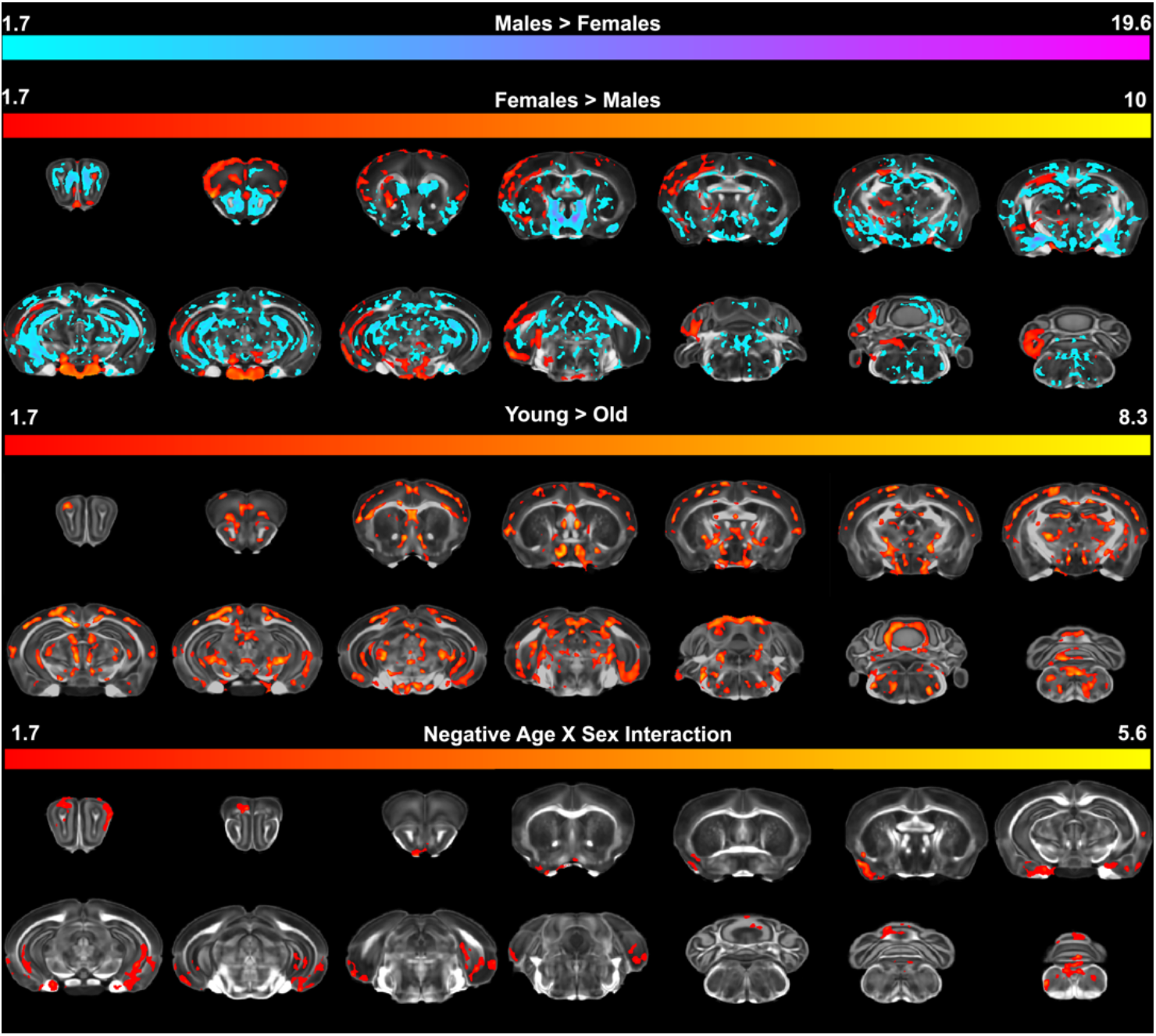
APOE2 targeted replacement mice showed sex and age specific morphometric differences, and the negative interaction of age by sex was indicative of faster changes in females. Males had larger clusters in the olfactory areas, dorsal caudate, piriform and cingulate cortices, preoptic areas, M1, amygdala, hippocampus, hypothalamus, visual cortex, and the left cerebellum (shown in blue). In contrast females had larger right caudate, hippocampus, entorhinal cortex, and cerebellum (shown in red). Relative to old mice, younger mice had larger somatosensory areas, preoptic, thalamic nuclei (zona incerta, medial geniculate), amygdala, ventral hippocampus, entorhinal cortices, medial pretectal, and visual cortex. The fornix was also larger in younger relative to older mice. There was a negative interaction indicating faster changes in females in olfactory areas, right piriform cortex, bilateral amygdala, ventral hippocampus and entorhinal cortex, and cerebellum. These results support increased vulnerability with aging and female sex in these areas. Increases in males are shown in blue, in females are shown in red. FDR corrected parametric maps show t and F values.

Relative to old mice, the younger mice had larger somatosensory areas, preoptic, thalamic nuclei (zona incerta, medial geniculate), amygdala, ventral hippocampus, entorhinal cortices, medial pretectal, and visual cortex. Interestingly the fornix was larger in younger mice relative to older mice. There was a negative age by sex interaction indicating faster changes in females in olfactory areas, right piriform cortex, bilateral amygdala, ventral hippocampus, entorhinal cortex, and cerebellum, denoting vulnerable regions with aging and female sex.

#### Diffusion Metrics

Regional based analyses indicated that the largest differences with age were located to the cingulate cortex, hypothalamus, posterior commissure, and lateral lemniscus, presubiculum, and primary somatosensory cortex (>30%) (**Supplementary Table 2**). No regional differences with sex survived FDR correction.

Voxel based fractional anisotropy analyses (**Figure 4**) indicated a midlife advantage for females, particularly for the frontal areas, cingulate cortex, insula, right ventral pallidum, dorsal thalamus, zona incerta, posterior hypothalamus, ventral hippocampus, amygdala, visual and entorhinal cortex, and cerebellum. Age effects were widespread for both gray and white matter, in particular the anterior commissure and corpus callosum (rostral part), but also the gray matter of the olfactory and premotor areas, hippocampus, thalamus, inferior and superior colliculi, brainstem and cerebellum. The negative interaction of age by sex suggests a faster decline in females, notably in the frontal association cortex, left motor cortex, insula, amygdala, ventral hippocampus and dentate gyrus, entorhinal cortex, periaqueductal gray, pons, and cerebellum.

**Figure 4.**
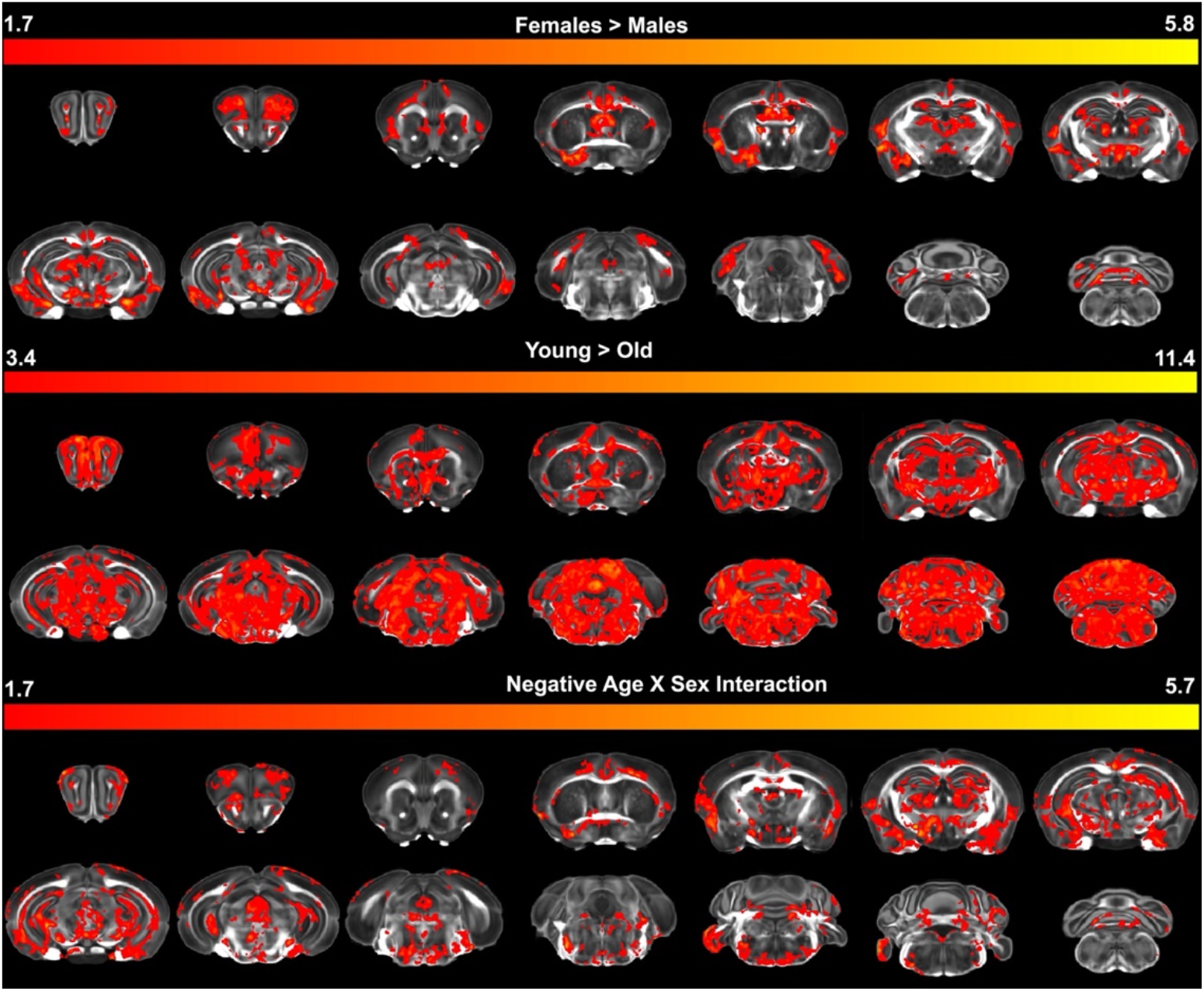
Diffusion tensor based FA estimates. Fractional anisotropy indicated an advantage for females, for the frontal areas, cingulate cortex, insula, right ventral pallidum, dorsal thalamus, zona incerta, posterior hypothalamus, ventral hippocampus, amygdala, visual and entorhinal cortex, and cerebellum. Age differences were widespread throughout the brain affecting both gray and white matter, in particular the anterior commissure, corpus callosum (rostral part), but also the gray matter of the olfactory and premotor areas, hippocampus, thalamus, inferior and superior colliculi, brainstem and cerebellum. There was a negative interaction of age by sex, suggesting a faster decline in females in the frontal association cortex, left motor cortex, insula, amygdala, ventral hippocampus and dentate gyrus, entorhinal cortex, periaqueductal gray and pons, and cerebellum. Increases in males are shown in blue, and in females in red. Parametric maps show t and F values.

#### SHORE /MAP Metrics

We estimated the return to origin probability (RTOP) (**Figure 5**), which indicated larger axonal diameters in males compared to females in the fimbria and fornix; as well as in small areas of the anterior corpus callosum, interior capsule and cerebral peduncles. The younger animals also had larger RTOP values in the corpus callosum, fimbria (septal region), and internal capsule. The age x sex interaction spared white matter.

**Figure 5.**
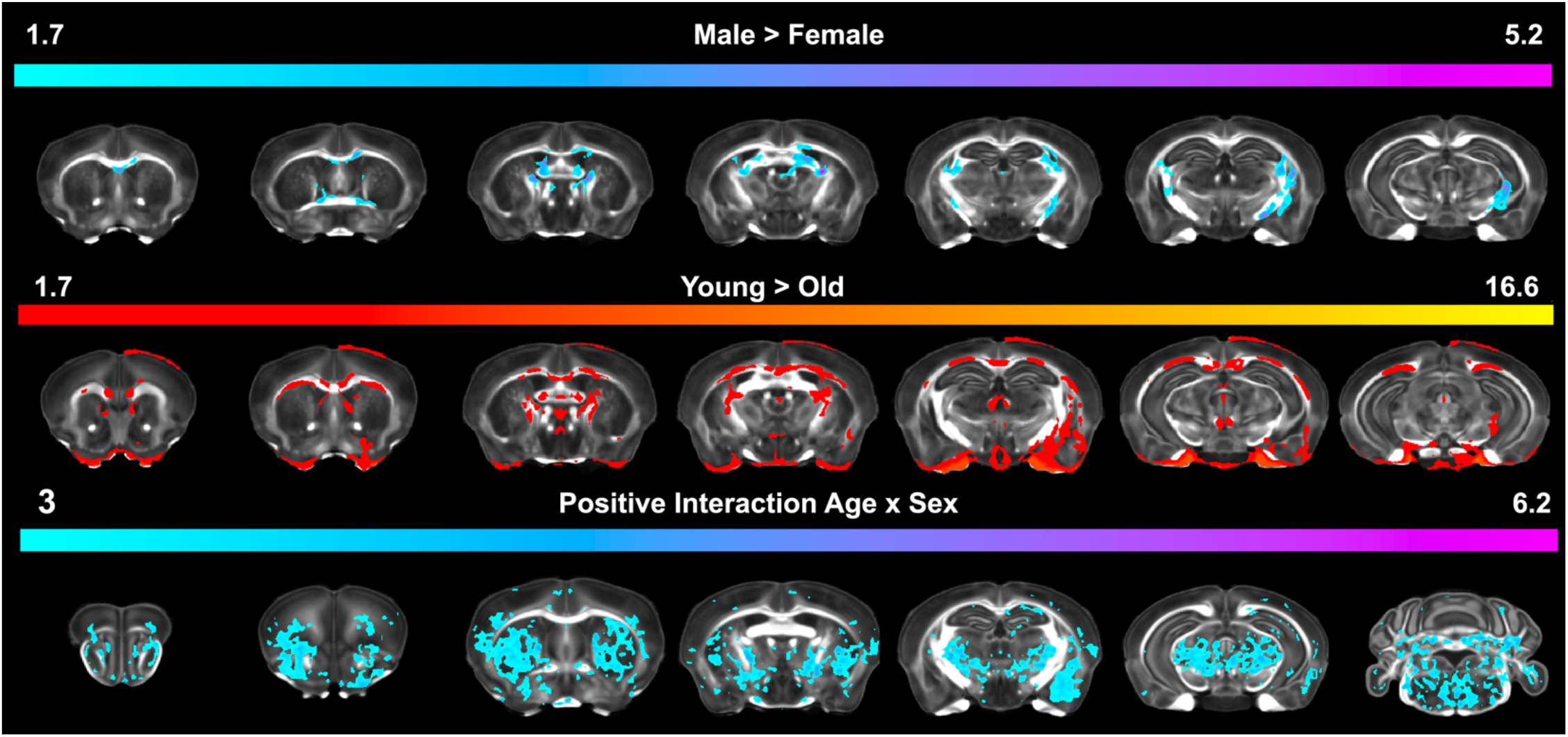
SHORE metrics like return to origin probability (RTOP) indicated increased cellularity, and axonal diameters in males versus females, and in younger versus older animals. There was a positive age x sex interaction for the gray matter areas, but not white matter, denoting faster increases in cellularity in males during aging and resilience of white matter tracts.

### Connectivity

#### Vertex Screening Based Detection of Structural Network Differences

The vertex screening method detected networks associated with age and sex separately (**Figure 6**), and these results supported a role of the thalamus, somatosensory cortices, and red nucleus that is involved in limb control, and motor cortex to cerebellar connections.

**Figure 6.**
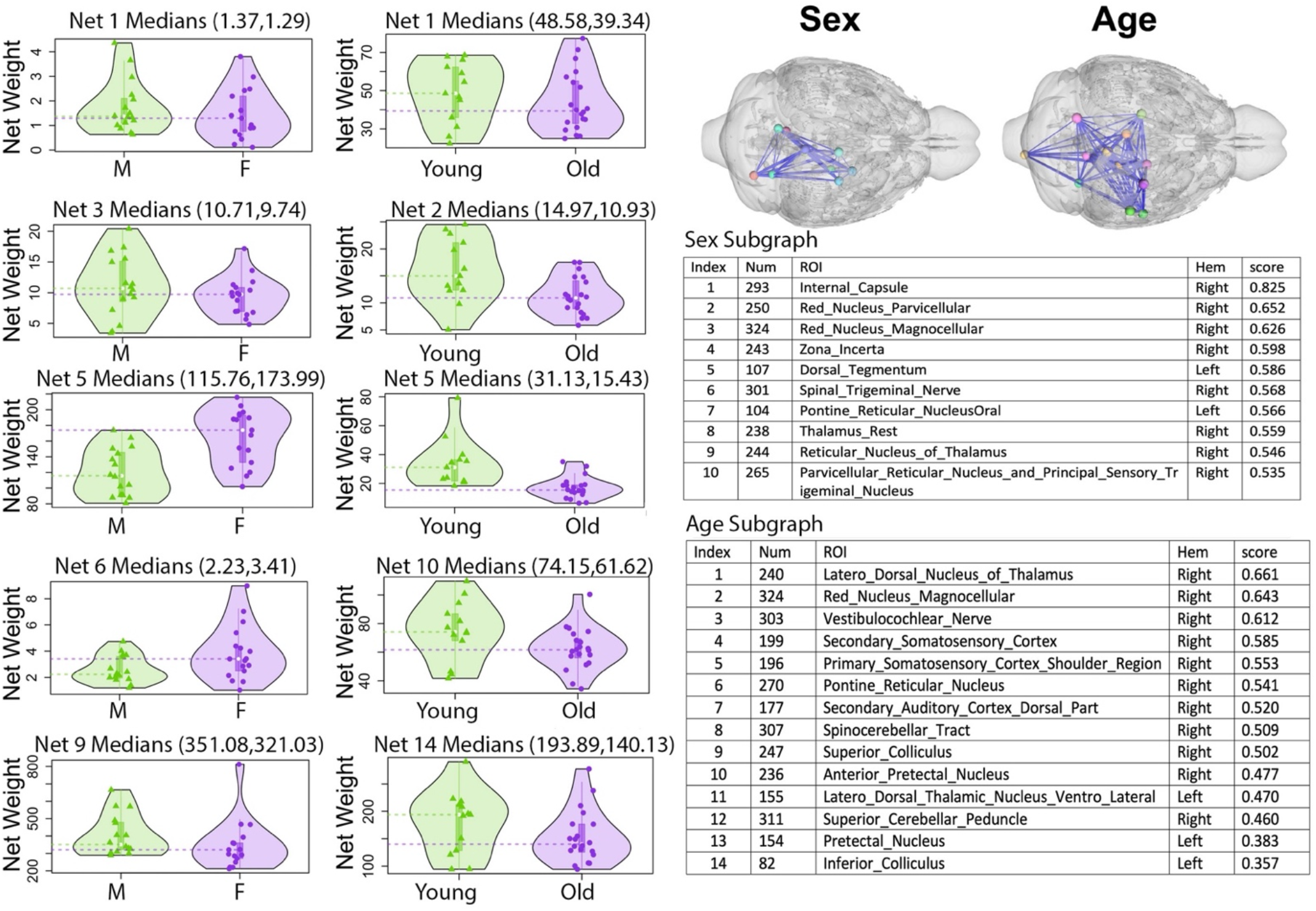
Vertex screening detected subgraphs that discriminate with age and sex. The left columns show sex differences. Net 1 (region 293): internal capsule; Net 3: red nucleus parvicellular part; Net 5: dorsal tegmentum; Net 6: spinal trigeminal nerve; Net 9: reticular nucleus of thalamus. The right column shows age differences for Net 1: laterodorsal thalamic nucleus; Net 2: red nucleus mangocellular; Net 5: primary somatosensory cortex; Net10: anterior pretectal nucleus; Net14: inferior colliculus. The spatial distributions of the sex and age vulnerable subgraphs are shown in the background of the anatomy, and the index in the atlas, hemisphere, and weights are indicated in the associated tables.

We validated the results through a series of Neural Network classifiers via a k=5-fold testing, and a 80:20 data split. The age category prediction accuracy estimated by a simple forward neural network (logistic) increased from 60% when using the whole connectome, to 78.5% when using the subgraph estimated through vertex screening. In contrast, sex prediction accuracy increased from 77.6% when using the whole connectome to 91.4% when using the subgraph estimated through vertex screening. When using 4 hidden layers in the neural network the accuracy for age category prediction increased from 72.3% when using the whole connectome, to 79.04% when using the subgraph estimated through vertex screening. The neural network with 4 hidden layers increased accuracy for sex prediction from 81.9% when using the whole connectome, to 94.28% when using the subgraph estimated through vertex screening. These results show that vertex screening significantly improved the accuracy of prediction over the full connectome.

The GCN prediction accuracy was 79% for both age and sex. Applying first vertex screening to connectomes resulted in only 10 regions being selected for age classification, and 14 regions for sex classification; accuracy increased to 82% for age, and to 85% for sex; while training time was reduced from 74 to 62 minutes for both age and sex. These results show that vertex screening significantly improved the accuracy of the prediction over the full connectome.

Sex differences in connectivity indicated a role for thalamic nuclei, including zona incerta, reticular nucleus, and the rest of the thalamus. Females had larger connectivity for the dorsal tegmentum (Net5), spinal trigeminal nerve (Net6), and lower connectivity in red nucleus and rest of thalamus, and the parvicellular reticular nuclei.

Age differences indicated decreases in the connectivity of the laterodorsal thalamic nucleus, red nucleus, primary somatosensory cortex, anterior pretectal nucleus, and inferior colliculus.

Both increases and decreases in connectivity were associated with age and sex differences, but changes were not uniform throughout the brain. The red nucleus was common to both age and sex vulnerable networks.

#### MCCA detection of vulnerable networks involved in learning and memory

MCCA detected brain networks associated with age and sex traits, and learning and memory (**Figure 7**), with a sum of the three correlations pairs of 1.68, and 95% bootstrap interval [0.80, 2.27]. The weights associated with the different traits were 0.99 for age, and 0.04 for sex; 0.04 for the normalized distance in the target quadrant and 0.99 for the absolute winding number, denoting a large relative importance of age in shaping brain networks, and for the winding number in describing memory and learning.

**Figure 7.**
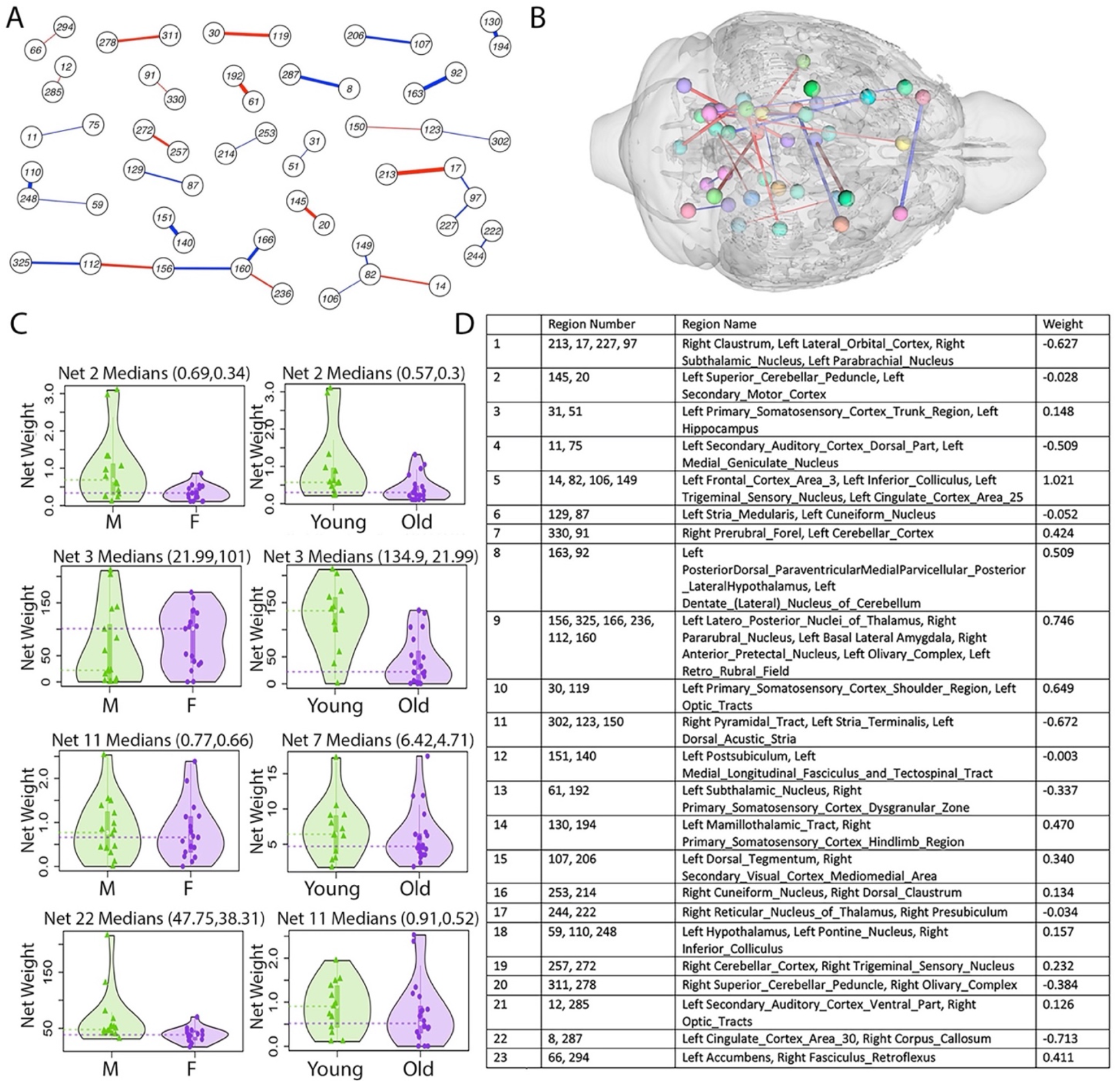
SMCCA detected brain networks associated with age, sex, and spatial memory as a composite risk factor. A. Positive weights are shown in blue, and negative weights are shown in red. B. 3D rendering of the brain subnetworks. C. Connectivity for selected subnetworks are shown using violin plots. D. The legend indicates the subgraphs components by index, region name, and their associated weight.

The largest subgraph weights were identified for a 4 node subgraph including the frontal area 3 FrA3, inferior colliculus, trigeminal sensory nucleus, and cingulate area 25 (weight = 1.02); followed by a 6 node subgraph including the latero posterior nucleus of the thalamus, pararubral nucleus, basal lateral amygdala, anterior pretectal nucleus, olivary complex, and the retrorubral field (weight = 0.75). The largest negative weight was associated with a pairwise connectivity of the cingulate cortex area 30 and the corpus callosum (−0.71), followed by a 3 node subgraph including the pyramidal tract, stria terminalis, and dorsal acoustic stria (−0.76).

Violin plots illustrated that some subgraphs were favored in males (e.g. for the secondary motor cortex and superior cerebellar peduncle), and other in females (e.g. hippocampus and primary somatosensory cortex, trunk region). Age effects were indicative of connectivity loss during aging (e.g. hippocampus and primary somatosensory cortex, trunk region).

#### Brain networks associated with peripheral /blood markers

Serum differential gene expression for the covariates age and sex are presented in **Supplementary Tables 3** and **4** respectively. Volcano plots emphasized genes showing differential expression with adjusted p < 0.05 for sex (**Figure 8A**) and age (**Figure 8B**). Applying sparse multi CCA (SMCCA) to the RNA-seq and connectome data effectively filtered age- and sex-related gene subsets (**Figures 8C and 8D, and Supplementary Tables 3 and 4)**associated with connectomes subgraphs and age (**Figure 9**), and respectively sex (**Figure 10**).

**Figure 8.**
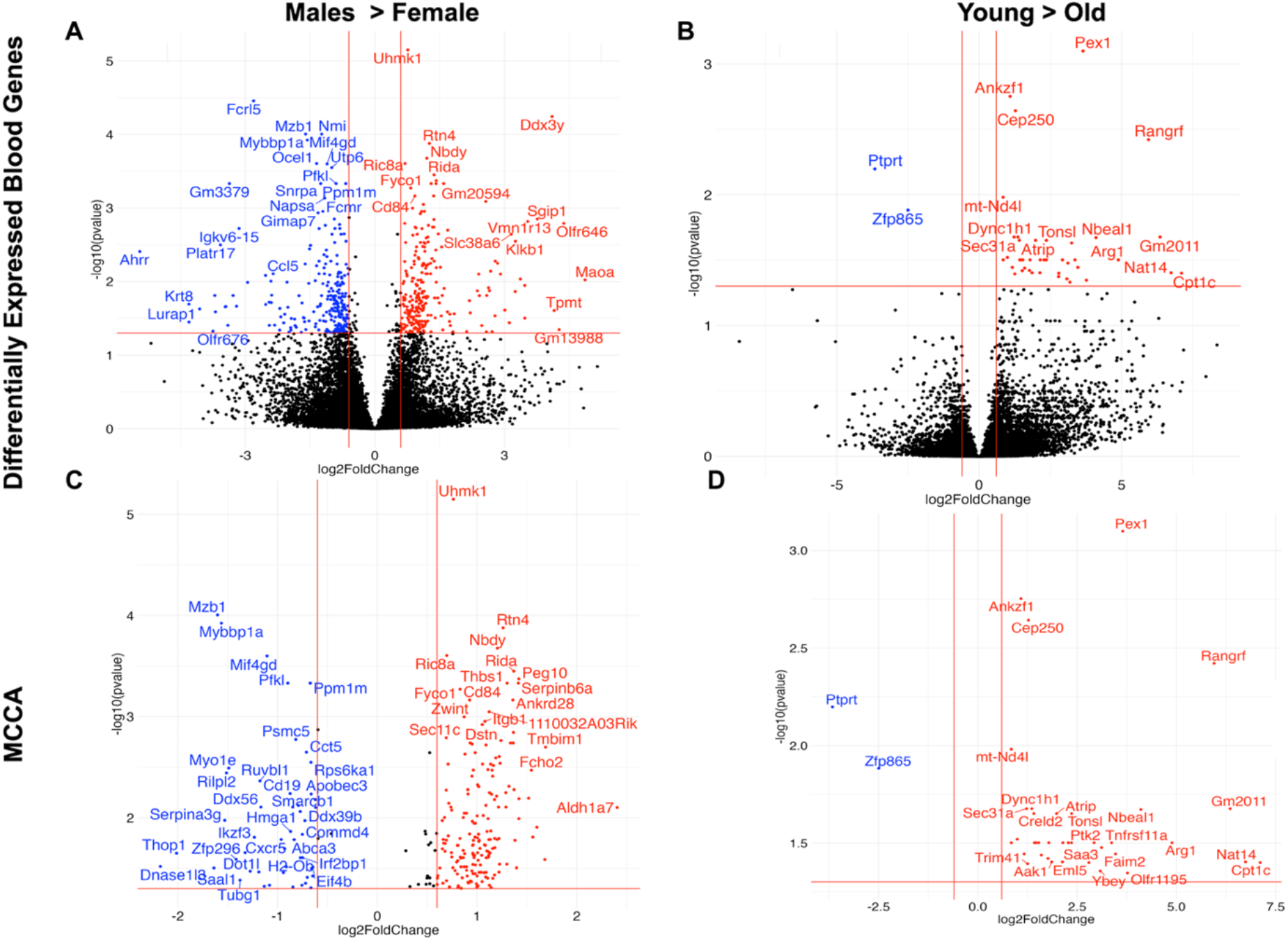
Differential gene expression for age and sex, before and after connectome-based filtering. The brain subgraphs for the SMCCA results and violin plots for distinct subnetworks connectivity are shown in **Figure 9**.

**Figure 9.**
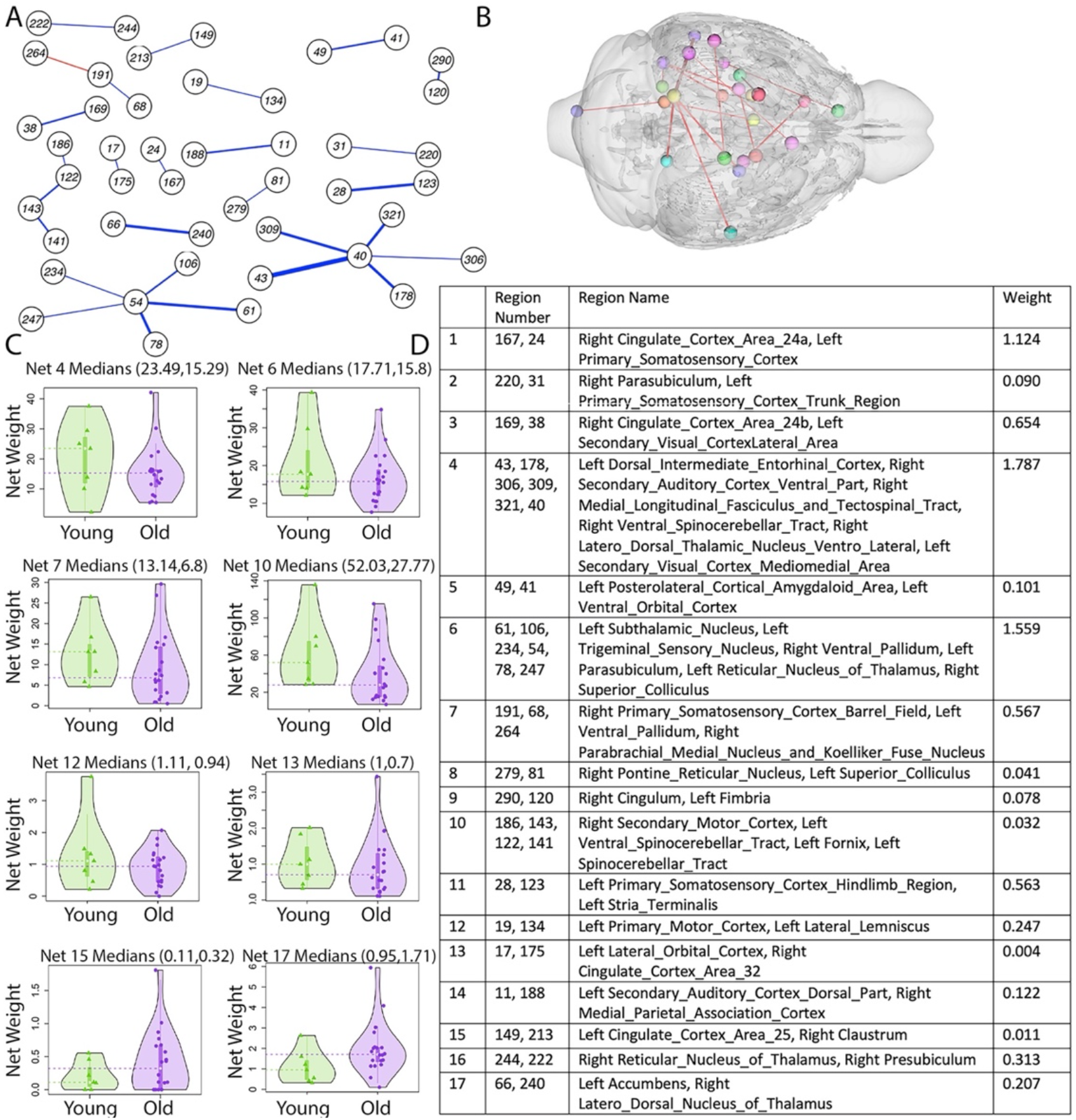
Brain networks associated with serum genes and aging in APOE2 mice indicate networks that decline and also age resilient or compensatory networks. A. Most subgraphs weights were positive (blue), however there was a negative weight (red) in the subgraph comprising the parabrachial nucleus Kolliker Fuse nucleus and the primary somatosensory cortex, the rest of the networks having positive weights. B. The spatial distribution of the subnetworks. C. Violin plots illustrated connectivity for selected subgraphs, for young and old mice. D. The highest weights were associated with Networks 4, 6, and 1 (according to rank order). These networks involved the entorhinal cortex, auditory cortex, latero dorsal thalamic nucleus, and visual cortex (Network 4); subthalamic nucleus, trigeminal sensory nucleus, ventral pallidum and parasubiculum, and superior colliculus (Network 6); the cingulate cortex area 24 and the primary somatosensory cortex (Network 1). The functional roles of these regions suggest changes in sensory (S1, V2), sensory-motor function (SC), and memory processing (entorhinal cortex, subiculum). We noted thalamic nuclei involved in spatial learning and memory (LDM), sensory processing, attention, cognition and transitioning between wakefulness and sleep (reticular nucleus (Li, Lopez-Huerta et al. 2020).

**Figure 10.**
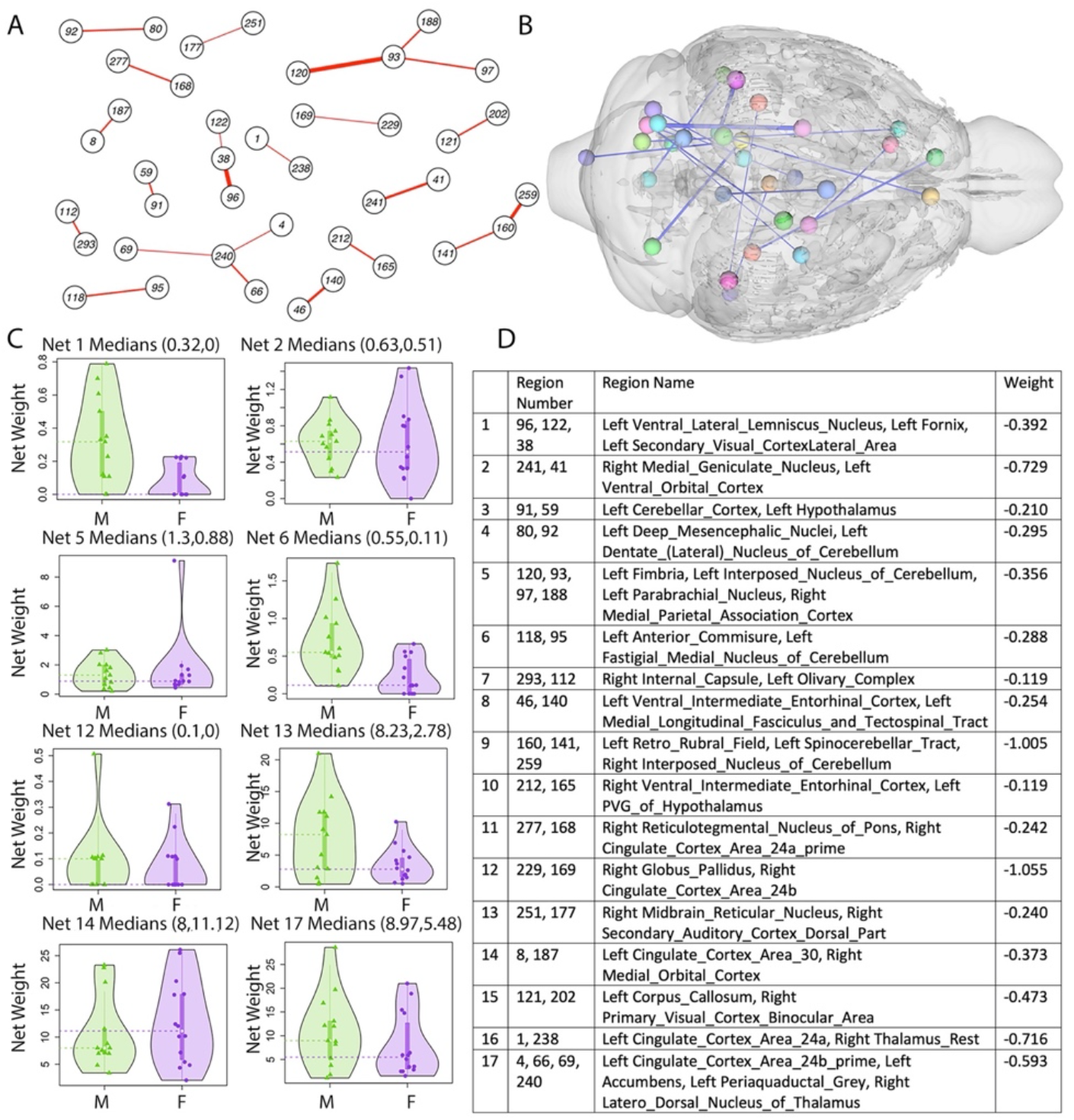
Brain networks associated with serum genes and sex in APOE2 mice. A. These subnetworks were characterized by negative weights, shown in red in the graph representation. B. The spatial distribution of the subnetworks. C. Violin plots showing connectivity for selected subgraphs, for males and females (M, F). D. The larges absolute wights were due to Networks 9, 2, 12, and 16. There regions involved the retrorubral field and interposed cerebellar nucleus (Network 9); medial geniculate nucleus, and ventral orbital cortex (Network 2); globus pallidus and cingulate cortex area 24b (Network 12); cingulate cortex area 24a and the rest of thalamus (Network 16).

After selecting the age-related genes the sum of 3 correlations between these age-related genes, connectomes, and traits (age, sex) was 2.48. The weight associated with sex was −0.04, and with age −0.99 The 95% bootstrapping confidence interval was [1.92, 2.71]. We tuned the hyperparameters so that most of the selected 42 genes (39) remained in the output.

After selecting the sex-related genes the sum of 3 correlations between these sex-related genes with connectomes and traits (age, sex) was 2.19. The weights associated with sex were 0.99, and for age 0.04. The 95% bootstrapping confidence interval was [1.86, 2.61]. We tuned hyperparameters to eliminate ~half of these 436 genes, resulting in 249 genes present in the output.

#### Age related blood transcriptome differences

Of the 15,934 genes, 44 showed an FDR-significant fold change ≤ 0.05. Of these, 42 showed positive fold changes with a range of log2FC of 0.8 to 7.1. Amongst the age-related genes surviving FDR correction, the largest FC was for Cpt1c (log2FC = 7.1, adjusted p = 0.04), involved with long chain fatty acids transport into mitochondria, and neuronal oxidative metabolism (Lee and Wolfgang 2012). Other genes that showed statistically-significant differential expression with large FC included Nat14 (log2FC = 6.7, SE = 1.7, adjusted p = 0.04), Rangrf (log2FC = 6.0, SE = 1.2, adjusted p = 0.004), Arg1 (log2FC = 4.9, SE = 1.2, adjusted p = 0.03), Nbeal1 (log2FC = 4.1, SE = 0.9, adjusted p = 0.02), and Olfr1195 (log2FC = 3.8, SE = 0.7, adjusted p = 0.001), Pex1 (log2FC = 3.6, SE = 0.7, adjusted p = 0.001), and Myo1e (log2FC = 1.8, SE = 0.4, adjusted p = 0.03). Most of these genes are present in the urea cycle, and also nervous processes (Olfr1195), cholesterol transport (Nbeal1), immune processes (Arg1), import of proteins into peroxisome (Pex1). Negative FC were observed for Ptprt (log2FC = −3.7, SE = 0.8, adjusted p = 0.006), and Zfp865 (log2FC = −2.5, SE = 0.6, adjusted p = 0.01). These genes are involved in signal transduction (Zfp865, Ptprt), and synapse organization (Ptprt).

GSEA for the 44 genes identified as FDR significant for differential expression by age identified the following annotations for GO terms or biological pathways (**Supplementary Table 5**): stress, signal transduction, transport, cellular nitrogen compound metabolic processes, immune system processes, and anatomical structure development.

After connectome based filtering (**Figure 9 D**) the genes that were also associated with connectomes and had the high absolute weights were Ankzfp1, with a role in cellular response to hydrogen peroxide and maintaining mitochondrial integrity under stress (van Haaften-Visser, Harakalova et al. 2017), as well as Pex1, Cep250, Nat14, Arg1, and Rangrf.

#### Sex related blood transcriptome differences

Of the 15,935 genes, 546 showed an FDR-significant fold change ≤ 0.05. Of these, 385 showed positive fold changes with a range of log2FC of 0.3 to 4.8 and 261 showed negative fold changes with a range of log2FC of −0.4 to −5.4.

Amongst the sex-related genes the largest FC were observed for Maoa (log2FC = 3.6, SE = 0.7, adjusted p = 0.0008) involved in catabolic processes, cellular nitrogen compound metabolic processes, and in the breakdown of the neurotransmitters: serotonin, epinephrine, norepinephrine, and dopamine. Several olfactory receptors (Olfr365, Olfr646) and thiopurine S-methyltransferase gene, TMPT (log2FC = 4.1, SE = 1.2, adjusted p = 0.02) also showed FDR-significant differences by sex. Negative FC were found for Ahrr (log2FC = −1.5, SE = 0.3, adjusted p = 0.003) which is involved in regulation of cell growth and differentiation; and Krt8 which plays a role in maintaining cellular structural integrity and also functions in signal transduction and cellular differentiation, Lurap1 which is involved in the positive regulation of cytokine production, and Oosp1 (Oocyte Secreted Protein 1) which may be involved in cell differentiation. Genes showing differential expression for sex and age included Myo1e (log2FC = −1.5, SE = 0.3, adjusted p = 0.003) and Ptprt (log2FC = 2.9, SE = 0.7, adjusted p = 0.008). GSEA for the 546 genes identified as FDR significant for differential expression by sex revealed the following annotations for GO terms or biological pathways (**Supplementary Table 6**): anatomical structure development (144 genes), transport (107 genes), cell differentiation (103 genes), response to stress and cellular nitrogen compound metabolic process (102 genes).

Connectome based filtering through MCCA for sex associated changes revealed (**Figure 9 C**) overexpressed genes, including Uhmk1 involved in phosphorylation of myelin basic protein and synapsin I, Serping6b, Fcho2, Peg10, and Ride.

There were four genes that were significant after FDR correction for both age and sex: Myo1e (log2FC = 31.5, SE = 0.3, adjusted p = 0.003), Ptprt (log2FC = 2.9, SE = 0.7, adjusted p = 0.008), Pex1 (log2FC = 3.6, SE = 0.7, adjusted p = 0.0008), and Creld2 (log2FC = 1.4, SE = 0.2, adjusted p = 0.02).

The connectome filtered genes differentially expressed with age or sex pointed to several enriched processes, with a role for biological and metabolic processes, cellular component organization and biogenesis, transport, and immune response, and also cell communication (**Figure 11**, **Supplementary Table 7**).

**Figure 11.**
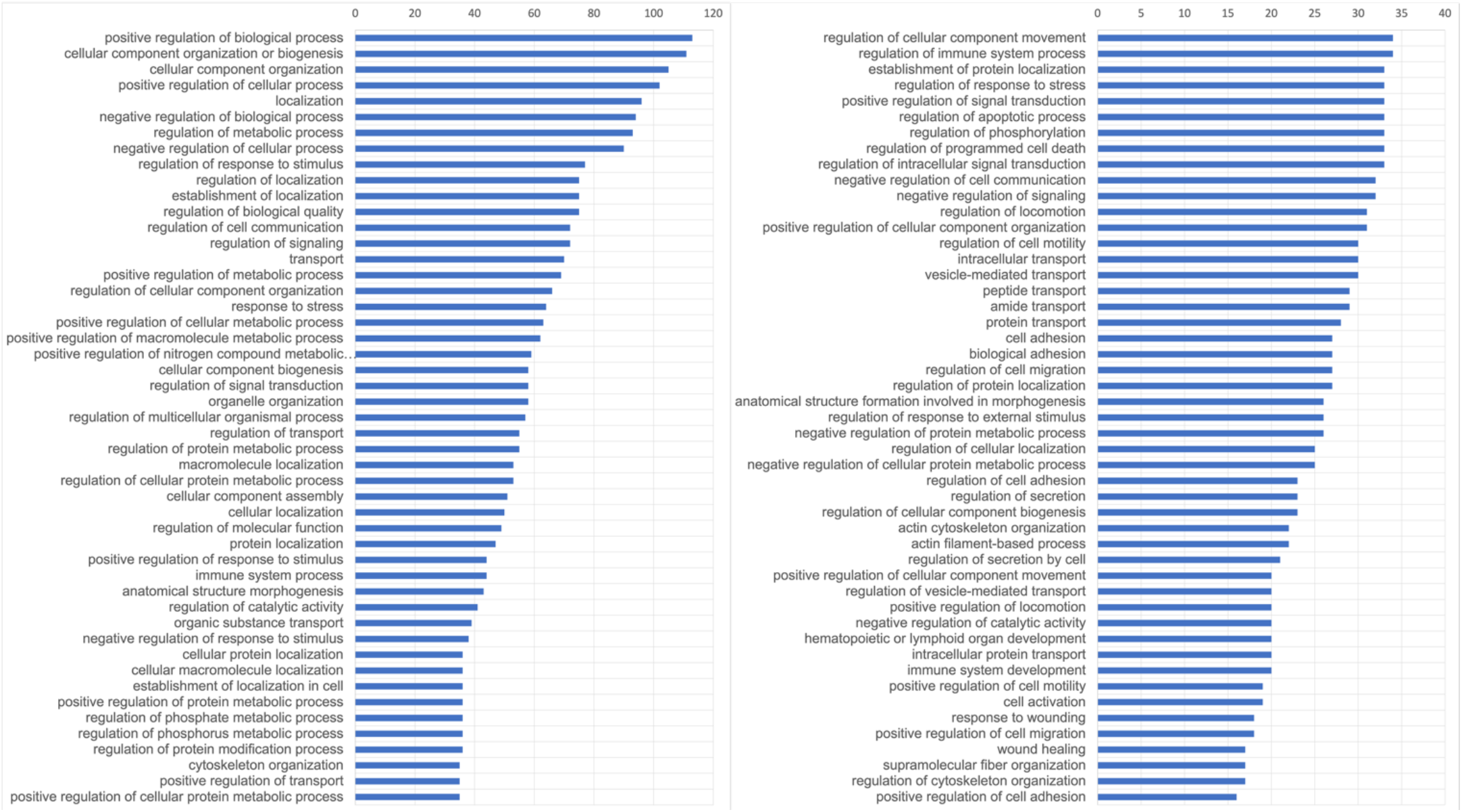
Enriched processes and pathways (n = 154) included regulation of cellular component organization or biogenesis (p = 2.1*10-7, N = 111 genes), localization (p = 1,2*10^-5^, N = 96 genes), metabolic process (p=0.04, N = 93 genes), response to stimulus (p = 5.4*10^-5^, N = 77 genes), signaling (p = 5.12, *10^-5^, N = 72 genes), transport (p = 7.6*10^-4^, N = 70 genes), response to stress (p = −0.001, N = 64 genes). Regulation of cell communication (p = 4.7*10^-5^, N = 72 genes) was highly significant.

Our results support that integrating blood based transcriptomics, imaging and traits (age, sex, cognition) help hone in on the mechanistic roles of APOE in modulating the transition from middle to old age, to help understand how APOE2 allele confer protection against pathological aging and AD, through maintaining cognition and brain connectivity via novel targets and pathways. Such modeling approaches might help devise strategies for enhancing the aging trajectory.

## Discussion

The APOE2 allele has been associated with longevity, protection against AD, and preserved cognition in AD, however it is not entirely benign, as it has also been associated with cerebral amyloid angiopathy, increased risk for PTSD, AMD, supranuclear palsy, and argyrophilic grain disease (Li, Shue et al. 2020). Our motivation stems from the relatively understudied role APOE2 plays to modulate synaptic integrity, brain connectivity, and response to inflammation and pathology that occur with aging, in particular during critical periods that mark the transition between middle and old age, or between an asymptomatic and symptomatic stage in neurodegenerative diseases. Given that female sex has been associated with vulnerability to age related cognitive decline and neurodegeneration, it is of interest to understand how APOE2 interacts with sex during aging. To help understand the role of APOE2 we investigated using humanized APOE2 targeted replacement mice how age and sex affected the transition from the human equivalent of middle age to old age. We have examined separately behavior, imaging and transcriptomic changes and identified joint variations in these domains using canonical correlation analyses.

We hypothesized that the APOE2 allele presence will result in only mild cognitive deficits with aging (Shinohara, Kanekiyo et al. 2016). Indeed, in our study, the Morris water maze learning trials revealed a significant interaction of age x stage (i.e. time) for the absolute winding number only. The probe tests performed 1 hour after the end of learning trials showed an effect of age both for the winding number and distance; while the probe administered 48 hours later revealed a significant effect of age for distance, percentage distance swam in the target quadrant, and the winding number. These results support that the winding number and spatial navigation strategies are sensitive to subtle cognitive changes in aging.

We expected little age related brain atrophy and sought to identify which brain regions were vulnerable or resilient during aging. The small but significant global brain atrophy (3%) due to aging was explained in part by atrophy in the hippocampal commissure, fornix, and cingulate cortex area 24 (~10%). Females had larger volumes relative to males for the bed nucleus of stria terminalis, subbrachial nucleus, postsubiculum (~10%), and claustrum (>5%), while males had larger volumes for the brachium of the superior colliculus, orbitofrontal cortex, frontal association cortex, and the longitudinal fasciculus of pons >9%). Age related atrophy was noted in white matter (anterior commissure, corpus callosum, etc.), as well as gray matter, in particular in olfactory cortex, frontal association area 3, thalamus, hippocampus and cerebellum. The amygdala was associated with both age and sex related vulnerability to aging, showing a faster decline in females. A negative age by sex interaction was also noted for the olfactory areas, right piriform cortex, bilateral amygdala, ventral hippocampus and entorhinal cortex, and cerebellum. We compare the VBA age associated atrophy patterns in APOE2 mice with (Shimada, Hosokawa et al. 1994) for a model of age-related brain atrophy. Aging induced in this SAM-P/10 mouse atrophy in the prefrontal cortex, piriform and entorhinal cortex, anterior olfactory nucleus, septum, amygdala, caudate-putamen, nucleus accumbens and cerebellar cortex. However, the hippocampus, diencephalon and brainstem structures showed no atrophy in the SAM-P/10. Our results support that changes with aging, and female associated vulnerability have both a common and also strain, i.e. genotype specific components.

Fractional anisotropy (FA) was used to characterize tissue microstructure, and showed brain wide changes with aging in white matter tracts including the anterior and posterior commissures, corpus callosum; and also in gray matter e.g. the olfactory areas, to septum, hippocampus, and cerebellum. Females had higher FA, in particular for frontal areas, cingulate cortex, insula, right ventral pallidum, dorsal thalamus, zona incerta, posterior hypothalamus, ventral hippocampus, amygdala, visual and entorhinal cortex, and cerebellum. However, a negative age by sex interaction was suggestive of faster decline in females for the frontal association cortex, left motor cortex, insula, amygdala, ventral hippocampus and dentate gyrus, entorhinal cortex, periaqueductal gray, and the cerebellum. Interestingly the interaction term spared largely white matter tracts in females during aging. We examined RTOP, which quantifies the probability of zero net displacements and reflects the presence of microscopic barriers and hindrances (e.g., cell membranes and filaments). RTOP was found to correlate (corr = 0.96) with axon diameter (Fick, Sepasian et al. 2017). In contrast to FA, RTOP showed localized differences with age and sex in corpus callosum, fimbria and septum/fornix. Faster changes were noted for males, but the interaction term indicated sparing of white matter tracts, which may be supportive of resilience of the brain networks in APOE2 males. The increase in estimated cellularity may be attributed to cellular swelling, increased numbers of activated microglia, or astrogliosis (Peters 2006, Wang, Wang et al. 2019), (Mattson and Arumugam 2018).

We used brain connectivity to reveal robust biomarkers for aging, sensitive to sex differences, since such estimates have the ability to integrate several markers for cellular density, axonal integrity, myelination, etc. Females had lower connectivity thalamus, but also in the red nucleus, and the parvicellular reticular nuclei. Aging led to decreased connectivity for the laterodorsal thalamic nucleus, red nucleus, primary somatosensory cortex, anterior pretectal nucleus, and inferior colliculus. The red nucleus was an indicator of vulnerability for both female sex and age, and suggest movement, tremors, and other limb related issues.

Our SCCA analyses were originally developed for building risk models identifying vulnerable subgraphs associated with composite risk traits in humans with APOE genotypes (Mahzarnia, Stout et al. 2022). Here we used similar integrative models over cognition and imaging in aging APOE2 mice and expanded the method to include transcriptomics. Our results revealed an important role for age (relative to sex) in shaping brain networks; meanwhile the winding number showed a good sensitivity to total distance and normalized distance in the target quadrant in particular for learning trials. Brain subgraph nodes that accounted for such changes included the frontal association cortex (involved in associative learning and memory formation (Nakayama, Baraki et al. 2015), cingulate area 25 (with a role in cardiovascular, endocrine and immune response (Alexander, Clarke et al. 2019), the latero posterior nucleus of the thalamus (involved in spatial learning and memory (van Groen, Kadish et al. 2002), anterior pretectal nucleus (visual system, somatosensory and pain sensation processing (Rees and Roberts 1993)), basal lateral amygdala (anxiety, reward, social behavior (Yang and Wang 2017)), and olivary complex (hearing, and also cerebellar motor learning (Welsh, Lang et al. 1995).

Whole blood based transcriptomics revealed that the largest positive fold change with aging was for Cpt1c, which regulates the beta-oxidation and transport of long-chain fatty acids into mitochondria, and has been shown to have a role in neuronal oxidative metabolism (Lee and Wolfgang 2012). Furthermore, GO annotations point to its role in lipid metabolic processes, transport, small molecule metabolic processes, and nitrogen compound metabolic processes. Amongst other genes differentially expressed with aging we identified Pex1, Myo1e, Pex1, and Arg 1. Pex 1 is involved with import of proteins into peroxisomes, peroxisome biogenesis and localization, ATP binding, protein binding. Myo1e encodes a nonmuscle class I myosin, whose proteins function as actin-based molecular motors, and may be involved in intracellular movement and membrane trafficking. Myo1e was found to be differentially expressed in disease associated microglia in mouse models of neurodegenerative disease (Sobue, Komine et al. 2021). Arg1 is a critical regulator of innate and adaptive immune responses. Negative FC were observed for Zfp865, involved in transcription coregulator activity, and Ptprt involved in signal transduction, cell adhesion and synapse formation, and cellular protein modification. We noted also PTK2 involved in integrin signaling, and Cep250, Olfr 646, 365 and 1195 involved with nervous system processes.

Out of 546 genes differentially expressed with sex, the largest positive FC was for Maoa, involved in catabolic processes, cellular nitrogen compound metabolic processes, and the breakdown of the neurotransmitters serotonin, epinephrine, norepinephrine, and dopamine. We noted significant positive FC for Olfr646 and Olfr365 involved in signal transduction and nervous system processes. In contrast the largest negative changes were noted for Ahrr, which plays a role in regulation of cell growth and differentiation; Krt8 which plays a role in maintaining cellular structural integrity, and SAAL1 is involved in response to proinflammatory stimuli.

Our study revealed a larger number of genes with significant expression changes with sex (546) than with aging (44), and associated with connectomes. For sex-based analyses the connectome based filtering through MCCA changes revealed a role for Uhmk1, Serping6b, Fcho2, Peg10, and Ride. These gene have been involved in anatomical structure development, sensory perception of sound, cell proliferation, differentiation and apoptosis, metabolism of amino acids. Genes that showed differential expression for sex in addition to age were Myo1e, Ptprt, Pex1 and Creld2. Several of these genes are involved in regulation of immune processes, efferocytosis, apoptosis and response to stress that have also been involved in the development of Alzheimer’s disease(Karch and Goate 2015, Neuner, Tcw et al. 2020, Romero-Molina, Garretti et al. 2022), although these studies have not clarified age- and sex-specific differences.

An important cellular component for these processes are microglia, macrophages of the brain parenchyma. Microglia play key roles in the maintenance and restoration of tissue homeostasis and the innate immune response(Romero-Molina, Garretti et al. 2022). Studies of microglial transcriptomics states in AD brains have identified specific disease-associated microglial (DAM) signatures where gene expression is markedly different from control brains. Myo1E is a gene found to be differentially-expressed in disease associated microglia in neurodegenerative disease mouse models and part of a DAM signature, specifically 1 of out 8 genes downregulated in AD precuneus (Sobue, Komine et al. 2021).

Other key neuronal molecular functions are also regulated by these differentially expressed genes identified in our study. Pex1 is part of the family of peroxin genes. These genes are involved with neuronal migration, myelination and brain development (Uzor, McCullough et al. 2020). Ptprt regulates synapse formation through interaction with cell adhesion molecules (Lee 2015). Creld2 augments protein folding and creates an interlink between the unfolded protein response axes through its interaction with proteins involved in the cellular stress response, promoting tolerance to endoplasmic reticulum stress and recovery from acute stress (Kern, Balzer et al. 2021).

An enrichment analysis revealed a role for biological and metabolic processes, cellular component organization and biogenesis, transport, and immune response. Of note, there is strong precedence for the involvement of pathways related to the regulation of the immune system, response to cellular stress and apoptotic processes from human genetic studies of AD (Karch, Cruchaga et al. 2014, Karch and Goate 2015, Neuner, Tcw et al. 2020, Romero-Molina, Garretti et al. 2022). Future studies are needed to explore the relationships amongst these findings to age and sex related differences in mice with all three humanized APOE major alleles.

Connectome maps related to serum genes during aging revealed vulnerable networks including the entorhinal cortex, auditory cortex, latero dorsal thalamic nucleus, and visual cortex; superior colliculus; the cingulate cortex area 24 and the primary somatosensory cortex. These findings suggest changes in sensory (S1, V1), and sensory-motor function (SC), and memory processing (entorhinal cortex, subiculum), including spatial learning and memory (LDM), sensory processing, attention, cognition and transitioning between wakefulness and sleep (reticular nucleus; (Li, Lopez-Huerta et al. 2020).

Our study has several limitations, the first one coming from a narrow age interval. However this age span covering the transition between middle and old age in mice was useful to generate several hypotheses about behavioral changes related to spatial navigation skills in particular, imaging markers pointing to a role for the cingulate cortices, entorhinal cortex and fornix for example, and genes that modulate response to stress and inflammation. Second, we chose mouse models with targeted replacement for human APOE genes, to provide a framework to understand human vulnerability to aging and sex differences, and these models have limitations in modeling human aging and female menopause associated changes. While APOE4 and APOE3 have been intensely studied, the APOE2 gene role in animal models of AD risk has been less studied. The human APOE2 allele is thought to favor longevity, to be protective against AD, and against cognitive decline in AD (Hua, Leow et al. 2008, Grothe, Villeneuve et al. 2017, Kim, Seo et al. 2017, Groot, Sudre et al. 2018), however there are limitations in using mouse models to understand how APOE2 modulates aging and disease mechanisms. In particular APOE2 mice develop type III hyperlipidemia, at much higher rates than humans (10 times more). Nevertheless, these mice have pure genetic backgrounds and are raised in well controlled environments, and thus can be used to understand aspects of aging, and reveal mechanisms and interactions that are difficult to understand directly in human populations with high heterogeneity (Zhao, Ren et al. 2020).

The judicious use of animal models can lead to a better understanding of the role of select gene co-expression networks in shaping the integrity of brain networks as a function of sex, age, and in particular the interaction of genetic and environmental factors, to determine the role of major APOE alleles in differentially modulating the risk for successful versus pathological aging and AD.

## Acknowledgement

We are grateful to Dr Nobuyo Maeda for the mouse donations, to NIH and the Bass Connections program for supporting our research. This work was supported by RF1 AG057895, R01 AG066184, U24 CA220245, RF1 AG070149, P30 AG072958.

## Figures List

**Supplementary Table 1. Volume**

**Supplementary Table 2. Fractional Anisotropy**

**Supplementary Table 3. Differentially expressed genes with age before and after connectome filtering Supplementary Table 4. Differentially expressed genes with sex before and after connectome filtering Supplementary Table 5. GO processes and gene networks associated with differentially expressed genes with age**

**Supplementary Table 6. GO processes and gene networks associated with differentially expressed genes with sex**

**Supplementary Table 7. GO processes and gene networks associated with connectome filtered genes**

